# Micro-DeMix: A mixture beta-multinomial model for investigating the fecal microbiome compositions

**DOI:** 10.1101/2023.12.12.571369

**Authors:** Ruoqian Liu, Yue Wang, Dan Cheng

## Abstract

Extensive research has uncovered the involvement of the human gut microbiome in various facets of human health, including metabolism, nutrition, physiology, and immune function. Researchers often study fecal microbiota as a proxy for understanding the gut microbiome. However, it has been demonstrated that this approach may not suffice to yield a comprehensive understanding of the entire gut microbial community. Emerging research is revealing the heterogeneity of the gut microbiome across different gastrointestinal (GI) locations in both composition and functions. While spatial metagenomics approach has been developed to address these variations in mice, limitations arise when applying it to human-subject research, primarily due to its invasive nature. With these restrictions, we introduce Micro-DeMix, a mixture beta-multinomial model that decomposes the fecal microbiome at compositional level to understand the heterogeneity of the gut microbiome across various GI locations and extract meaningful insights about the biodiversity of the gut microbiome. Moreover, Micro-DeMix facilitates the discovery of differentially abundant microbes between GI regions through a hypothesis testing framework. We utilize the Inflammatory Bowel Disease (IBD) data from the NIH Integrative Human Microbiome Project to demonstrate the effectiveness and efficiency of the proposed Micro-DeMix.

**Key MessagesKey Messages:**

- Micro-DeMix is a computational tool for understanding the heterogeneity of the gut microbiome across GI locations.
- Micro-DeMix facilitates the detection of differentially abundant microbes along the GI tract.
- Micro-DeMix detects that in IBD populations, the lower GI tract exhibits a larger abundance of Firmicutes and Bacteroidetes, whereas the upper GI tract is predominated by Proteobacteria and Firmicutes.

## 1 Introduction

The human gut microbiome and its role in host health have been the subject of extensive research, establishing its involvement in human metabolism [7], nutrition [34], physiology [2], and immune function [33, 4]. Recent studies have also demonstrated the association between the gut microbiota and the emergence of obesity [3], metabolic syndrome and the onset of type 2 diabetes [17, 7, 18]. Moreover, altered composition and function of the gut microbiota has been found associated with chronic diseases [28] ranging from gastrointestinal inflammatory [30] and metabolic conditions to neurological [6], cardiovascular [32], and respiratory illnesses [9, 5].

Researchers often study fecal microbiota as a proxy for understanding the gut microbiome since fecal samples are noninvasive and easy to collect [23]. However, the fecal microbiome alone may not provide a complete picture of the entire gut microbial community, because it ignores the heterogeneity of the gut microbiome across gastrointestinal (GI) locations [1, 22]. Indeed, recent studies have revealed significant variations in microbial composition and functions along distinct segments of the small intestine [21]. Furthermore, in the small intestine, the families Lactobacillaceae and Enterobacteriaceae dominate, whereas the colon is characterized by the presence of species from the families Bacteroidaceae, Prevotellaceae, Rikenellaceae, Lachnospiraceae, and Ruminococcaceae [8]. In addition, researchers applied the spatial metagenomics approach to the mouse colonic microbiome and discovered a heterogeneous distribution of taxa in the gut [31]. However, this spatial metagenomics approach has its limitations in human-subject research due to its invasive nature. Therefore, there is a pressing need to develop computational methods that use existing fecal samples to understand the heterogeneity of the gut microbiome across various GI locations and to extract meaningful insights about the biodiversity of the gut microbiome.

We bridge this gap by developing Micro-DeMix, a novel statistical model for deconvoluting the fecal microbiome with respect to a reference microbiome data set collected at a specific GI location. Our model is motivated by the data derived from the Inflammatory Bowel Disease (IBD) Multiomics Database (IBDMDB) project from the NIH Integrative Human Microbiome Project (iHMP), which followed 132 individuals and collected 1,785 stool samples and 651 intestinal biopsies over time [15]. This integrated study, thus, motivated the development of Micro-DeMix which can effectively integrate the microbial profiles from diverse gastrointestinal locations and those from the stool samples.

Micro-DeMix is a mixture beta-multinomial model that can quantify the composition of fecal microbiomes. The “multinomial part” of Micro-DeMix effectively handles the compositional nature of microbiome data by directly focusing on the baterial proportions rather than counts. The “beta part” further adjusts for the sample heterogeneity that may impact the composition of fecal microbiomes by incorporating important demographic and clinical variables. Moreover, Micro-DeMix facilitates the discovery of differentially abundant microbes between GI regions through a theoretically justified and computationally efficient hypothesis testing framework.

The rest of the paper is organized as follows. In Section 2, we describe the Micro-DeMix framework, develop a maximum-likelihood-based estimation procedure, and propose a hypothesis-testing framework for detecting differentially abundant microbes between GI locations. In Section 3, we conduct simulation studies to demonstrate the effectiveness of Micro-DeMix in terms of the estimation accuracy and the power of the hypothesis testing method. In Section 4, we apply our method to the Inflammatory Bowel Disease (IBD) microbiome data from iHMP to elucidate the composition of fecal microbiome in IBD patients. We close with a discussion of our method in Section 5.

## 2 Methods

### 2.1 Model

In this section, we present Micro-DeMix, a beta-multinomial framework for modeling microbial abundance in stool samples. Our model considers a multivariate extension of the beta-binomial model proposed in a recent research [24] which models individual taxa separately. As shown later, this multivariate extension brings several statistical and computational challenges due to an additional sum-to-one constraint.

Let *y*_*ig*_ denote the observed absolute abundance for taxon *g* in the *i*-th stool samples for *g* = 1, …, *G* and *i* = 1, …, *S*. We model each **y**_*i*_ = (*y*_*i*1_, …, *y*_*iG*_)^⊤^ using a multinomial model

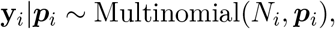

where 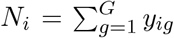 and ***p***_*i*_ = (*p*_*i*1_, …, *p*_*iG*_)^⊤^ is the underlying true microbial proportions. We are particularly interested in how the stool microbiome differs from the rectum microbiome, given the comprehensive data available for the rectum from the iHMP IBD study and the fact that the rectum is the final part of the large intestine. While we focus on the rectum due to both data availability and biological relevance, the methodology can seamlessly extend to other segments of the gastrointestinal tract, including the ileum, jejunum, colon, and beyond.

Acknowledging that the stool microbiome consists of microbes from rectum and other GI locations (mostly, the small intestine), we assume

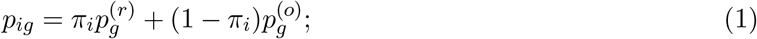

here, *π*_*i*_ *∈* (0, 1) denotes the latent proportion of the rectal microbiome in the *i*-th stool sample; 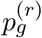 and 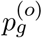 denote the proportion of taxon *g* in the rectum and other GI locations, respectively. In addition, we let 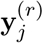 denote the reference rectum microbiome data for the *j*-th subject for *j* = 1, …, *S*^(*r*)^.

To allow *π*_*i*_ to vary across individuals, we assume

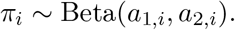

Hence, we write the joint density function of (*y*_*ig*_, *π*_*i*_) as

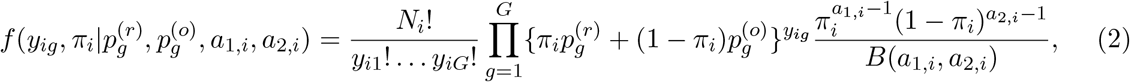

where *B*(*a*_1,*i*_, *a*_2,*i*_) = Γ(*a*_1,*i*_)Γ(*a*_2,*i*_)*/*Γ(*a*_1,*i*_ + *a*_2,*i*_). Since *π*_*i*_ is not observed, we integrate out *π* and obtain the density function of *y*_*ig*_:

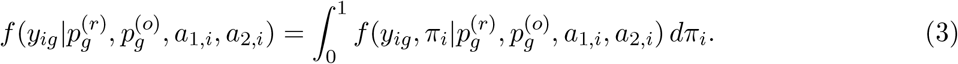

To capture the association between *π*_*i*_ and the subjects’ characteristics, we further link each *π*_*i*_ with covariates **x**_*i*_ through a beta-regression model. With additional reparameterization

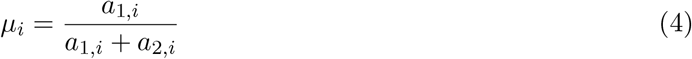

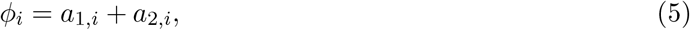

we consider

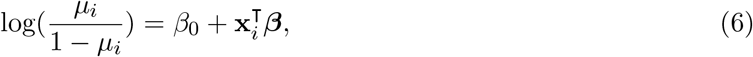

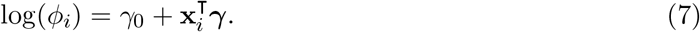

Some calculations yield that *µ*_*i*_ is the expected proportion of the rectum microbiome in the *i*-th stool sample, that is, *µ*_*i*_ = 𝔼(*π*_*i*_), and Var(*π*_*i*_) = *µ*_*i*_(1 − *µ*_*i*_)*/*(1 + *ϕ*_*i*_), where *ϕ*_*i*_ known as the precision parameter.

Thus, the log-likelihood of observing *{y*_*ig*_ : *i* = 1, …, *S*; *g* = 1, …, *G* in stool samples is

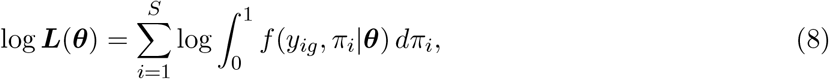

where 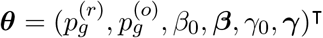.

**Remark 1**. *Under the proposed model, we derive the mean and variance of y*_*ig*_:

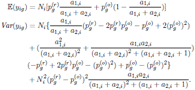

*See proof in Appendix*.

### 2.2 Estimation

We estimate ***θ*** by maximizing the log-likelihood function (8). We first estimate 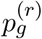 using the external rectum microbiome data for *g* = 1, …, *G*. Recall that 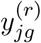 denotes the observed absolute abundance for taxon *g* and sample *j, j* = 1, …, *S*^(*r*)^, collected from rectum. Let 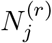 denote the sequencing depth for the *j*-th sample from rectum. We assume the absolute abundance to follow a multinomial distribution

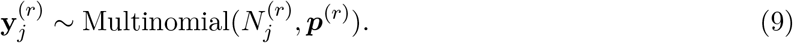

Based on this model, we estimate 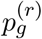 with the sample proportions

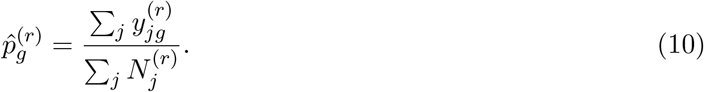

After replacing 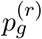 in (8) by 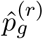, we estimate the remaining parameters using an iterative procedure with the main steps given in Algorithm 1.

Specifically, in the *m*-th iteration, we obtain 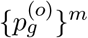 by maximizing the objective function (8) given *{β*_0_, ***β***, *γ*_0_, ***γ****}*^*m−*1^. Since the integral in (8) does not have a close form, we used the Gauss-Legendre

#### Algorithm 1

Estimating *{*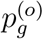, *β*_0_, ***β***, *γ*_0_, ***γ****}* for Micro-Demix

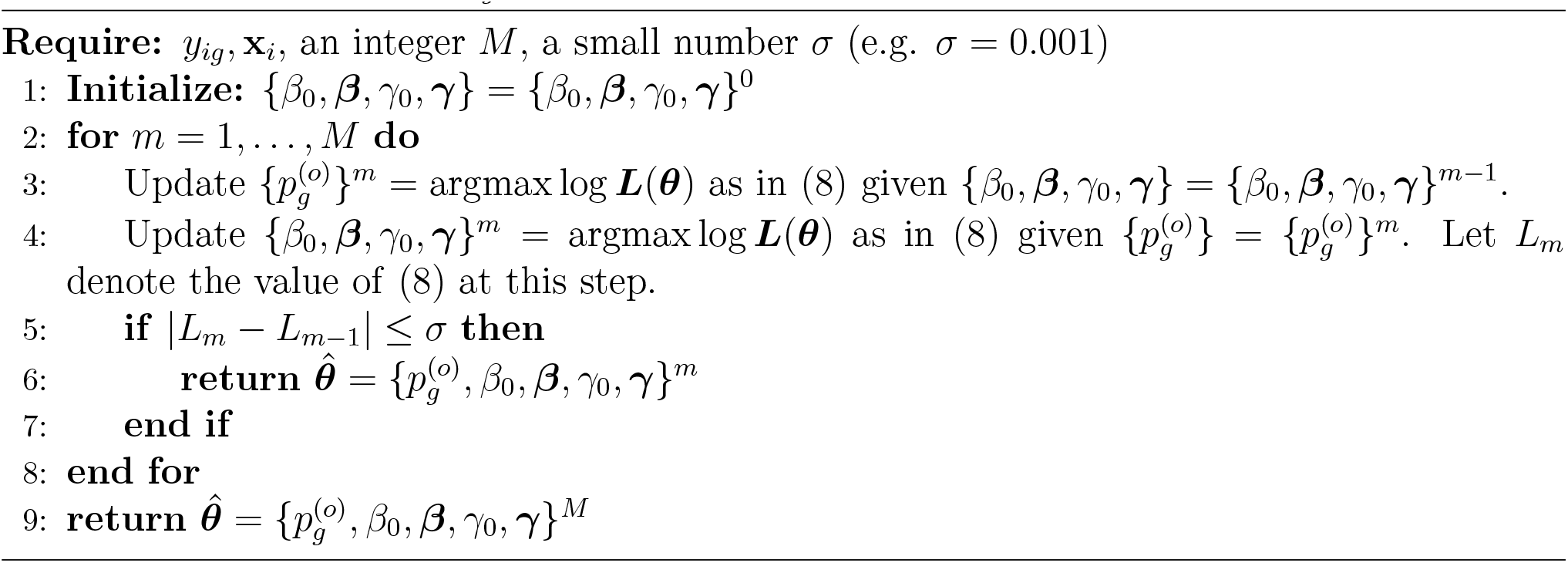

quadrature [14] to approximate it. Solving for 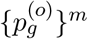 is a constrained optimization problem because 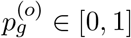 and 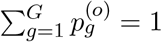. To address this problem, we consider the parameterization

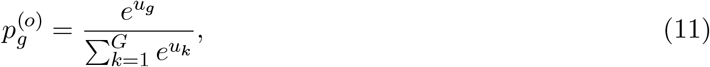

where *u*_*g*_ *∈* ℝ for *g* = 1, …, *G*. With this re-parametrization, we first obtain *{u*_*g*_*}*^*m*^ using the Nelder-Mead optimization algorithm from R package optimx and then calcualte 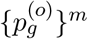 from *{u*_*g*_*}*^*m*^ using (11). Finally, given 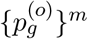, we obtain the optimal set *{β*, ***β***, *γ*, ***γ****}*^*m*^ using the Nelder-Mead optimization algorithm by maximizing log-likelihood function (8). We repeat this iterative procedure until we achieve convergence or reach the maximum number of iterations.

### 2.3 Hypothesis Testing

The proposed Micro-DeMix framework also facilitates the investigation of the biodiversity of the fecal microbiome by detecting microbial groups that are differentially abundant in rectum and other GI locations. Specifically, we consider a hypothesis testing problem with *H*_0_ : ***p***^(*o*)^ = ***p***^(*r*)^, where ***p***^(*o*)^ and ***p***^(*r*)^ are introduced in Section 2.1, representing the true proportions of the microbes of interest in the rectum and other GI locations, respectively.

Under *H*_0_, we have ∥***p***^(*o*)^ *−* ***p***^(*r*)^∥_2_ = 0, motivating the following test statistic:

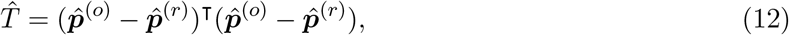

where 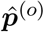 are maximum likelihood estimators obtained from Algorithm 1 in Section 2.2, and 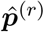 is given in (10). To figure out the null distribution of 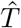, we first rewrite the test statistic 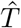:

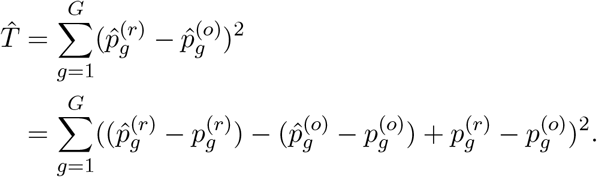

Under *H*_0_, we get

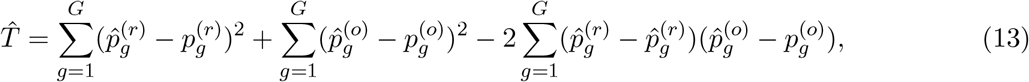

indicating that 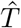 is a function of 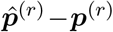 ***p***^(*r*)^ and 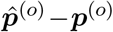 ***p***^(*o*)^ under *H*_0_. Thus, finding the null distribution of 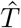 reduces to finding the null distributions of 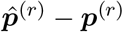 ***p***^(*r*)^ and 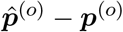. Specifically, applying the central limit theorem based on model (9), we have

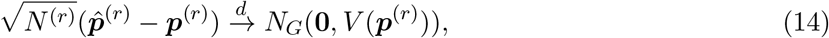

where

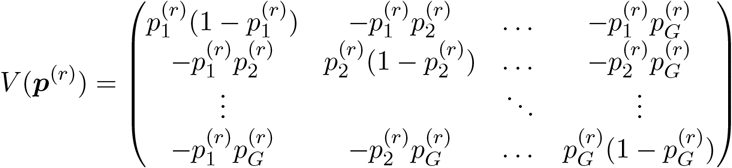

and 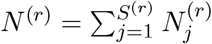. Also, under *H*_0_, the joint density function in (2) reduces to

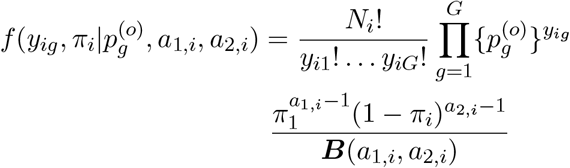

and the density function (3) becomes

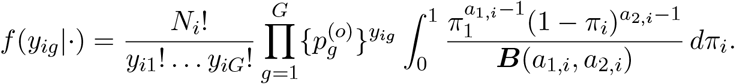

As the integral of the beta density is 1, we have

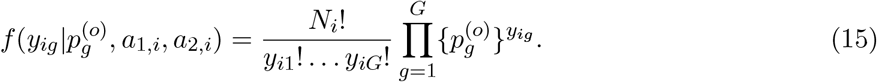

This indicates that under *H*_0_, the proposed Beta-multinomial model in Section 2.1 reduces to a multinomial model. Thus, under *H*_0_, the MLE derived from the mixture beta-multinomial model is equivalent to that derived from the multinomical model. Thus, we can apply the CLT again and obtain

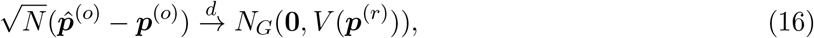

where 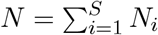 is the total counts of all samples collected from stool.

Assuming no overlapping between stool samples and rectum samples, we establish the asymptotic distribution of 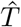 in the following result.

#### Proposition 1

*Let random vector* (***y***_1_, ***y***_2_) *follow the multivariate normal distribution*

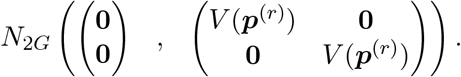

*Suppose* ***p***^(*o*)^ = ***p***^(*r*)^ *and N* ^(*r*)^*/N → K >* 0. *Then* 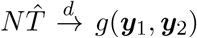 *as N → ∞, where* 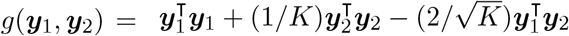

As a special case of Proposition 1, when *K* = 1 or *N* ^(*r*)^ = *N*, we have 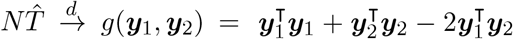. Based on Proposition 1, we develop the following simulation-based algorithm for calculating the p-value.

#### Algorithm 2

Simulation method of testing *H*_0_ : ***p***^(*o*)^ = ***p***^(*r*)^

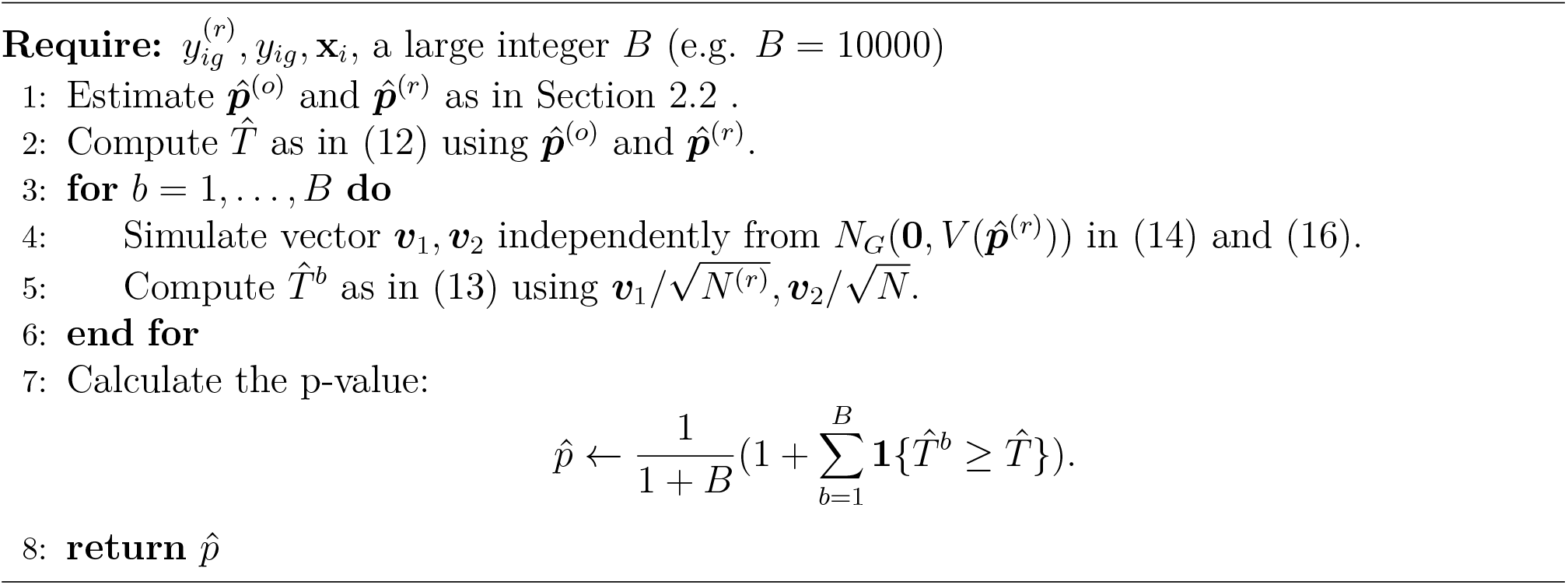

## 3 Simulation Study

We now investigate the finite-sample performance of Micro-DeMix using simulation. We studied the accuracy in the estimation of ***p***^(*o*)^ and the type I error rate and power when testing for differential abundance between the gut microbiome in the rectum and other GI locations.

For rectum samples, we simulated the microbial counts 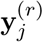 from a Multinomial(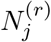, ***p***^(*r*)^) distribution with 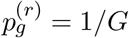 for *g* = 1, …*G* and *j* = 1, …, *S*^(*r*)^. We next generated the stool microbiome data using the proposed beta-multinomial model. Specifically, we first set (*β*_0_, *β, γ*_0_, *γ*) = (0.1, 0.1, 0.2, 0.1) and generated the covariate *x*_*i*_ from the standard normal distribution. We then randomly generated *π*_*i*_ from Beta(*a*_1,*i*_, *a*_2,*i*_), where *a*_1,*i*_ and *a*_2,*i*_ are defined in (4) - (7). We assigned *u*_*g*_ at evenly spaced values from -1 to 1 for *g* = 1, …, *G−* 1 and let 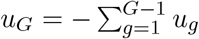. We computed 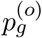 from *u*_*g*_ according to (11). Finally, we simulated the stool microbiome data **y**_*i*_ from a Multinomial(*N*_*i*_, ***p***_*i*_) distribution where ***p***_*i*_ is defined in (1) for *i* = 1, …, *S*.

### 3.1 Estimation

In this subsection, we assess the performance of Micro-DeMix in estimating ***p***^(*o*)^ under six settings. We considered *S* = *S*^(*r*)^ = 100, 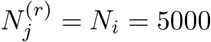, 10000, 15000 for all *i* and *j*, and *G* = 5, 10. We reported relative squared errors (RSE) to quantify the discrepancy between our Micro-DeMix estimators 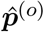 and the true parameters according to

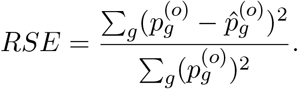

The mean and standard deviation of RSE under all six settings are presented in Table 1. It was observed that as the library size increases, both the mean and standard deviation of the RSE tend to decrease. This indicates that the accuracy of estimation improves as we increase the library size. Furthermore, it is noticeable that the RSE’s for G=10 are greater than those for G=5. This phenomenon occurs because the true parameters diminish as the total number of taxa increases. Specifically, by design, 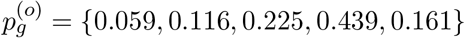 when *G* = 5, and 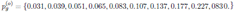 when *G* = 10. Due to the sum-to-one constraint, the true signal tends to decrease when G increase. This makes estimation more challenging and contributes to a decrease in relative accuracy.

**Table 1:** Mean and Standard Deviation of RSE obtained under each setting while estimating ***p***(*o*) in Micro-Demix.

| Library Size | G = 5 |  |  | G = 10 |  |  |
| --- | --- | --- | --- | --- | --- | --- |
|  | 500 | 1000 | 5000 | 500 | 1000 | 5000 |
| Mean | 0.00049 | 0.00035 | 0.00014 | 0.00557 | 0.00501 | 0.00364 |
| Stand Dev | 0.00045 | 0.00035 | 0.00027 | 0.00261 | 0.00201 | 0.00111 |

### 3.2 Type I Error Rate and Power

In this subsection, we assess the empirical type-I error rate and power of the proposed Micro-DeMix testing procedure to detect differential abundance between rectal microbes and microbes from other GI locations.

The data-generating process was the same as that in the previous section except for a few modifications to facilitate the assessment of the type-I error rate and power. Specifically, we set (*β*_0_,*β,γ*_0_,*γ*) = (0.1, 0.1, 0.7, 0.2). For each *G*, we generated ***p***^(*r*)^ by letting each 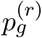 and 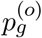 differ by 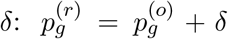 for *g* = 1, …, *G/*2 and 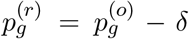 for *g* = *G/*2 + 1, …, *G*. This generating process ensures 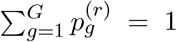 for any *δ*. We considered relatively weak signals, i.e., *δ* = 0, 0.0002, 0.0004, …, 0.002, for *G* = 10, and further decreased the signal strength by considering *δ* = 0, 0.0001, 0.0002, …, 0.0001 for *G* = 20. In both settings, we considered 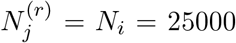 to ensure a large enough library size for signal detection.

We implemented the proposed Micro-DeMix testing procedure (detailed in Algorithm 2) to produce a p-value for testing *H*_0_ : ***p***^(*r*)^ = ***p***^(*o*)^ based on 10000 simulated data sets from the null distribution; that is, *B* = 10, 000 in Algorithm 2. Under the significance level *α* = 0.05, we examined the type-I error rate and power of the Micro-DeMix test for *δ* = 0 and *δ >* 0, respectively, based on 100 independent replications.

As a comparison, we considered an alternative testing procedure based on the asymptotic normality shown in (16). Specifically, under *H*_0_, the Micro-DeMix estimator of ***p***^(*o*)^ reduces to the MLE of ***p***^(*o*)^ based on (15), denoted by 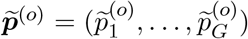 with 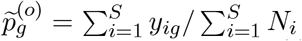 for *g* = 1, …, *G*. Therefore, one could generate an alternative p-value for testing *H*_0_ by replacing 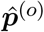 with 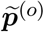 Algorithm 2. Since this alternative p-value is derived from the central limit theorem of the Multinomial distribution, we refer to it as multiCLT hereafter.

The results are shown in Figure 1. The value *t* on x-axis represents the total signal, and some calcalations yield that 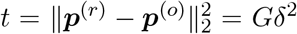. The points at 0 on x-axis representing the scenario that ***p***^(*r*)^ = ***p***^(*o*)^. For *G* = 10, the type-I error is 0.036 for MultiCLT and 0.063 for Micro-DeMix. It appears that MultiCLT is more conservative, while Micro-DeMix has a slight type-I error inflation. For *G* = 20, both methods demonstrate a comparable type-I error rate of 0.063.

**Figure 1:**
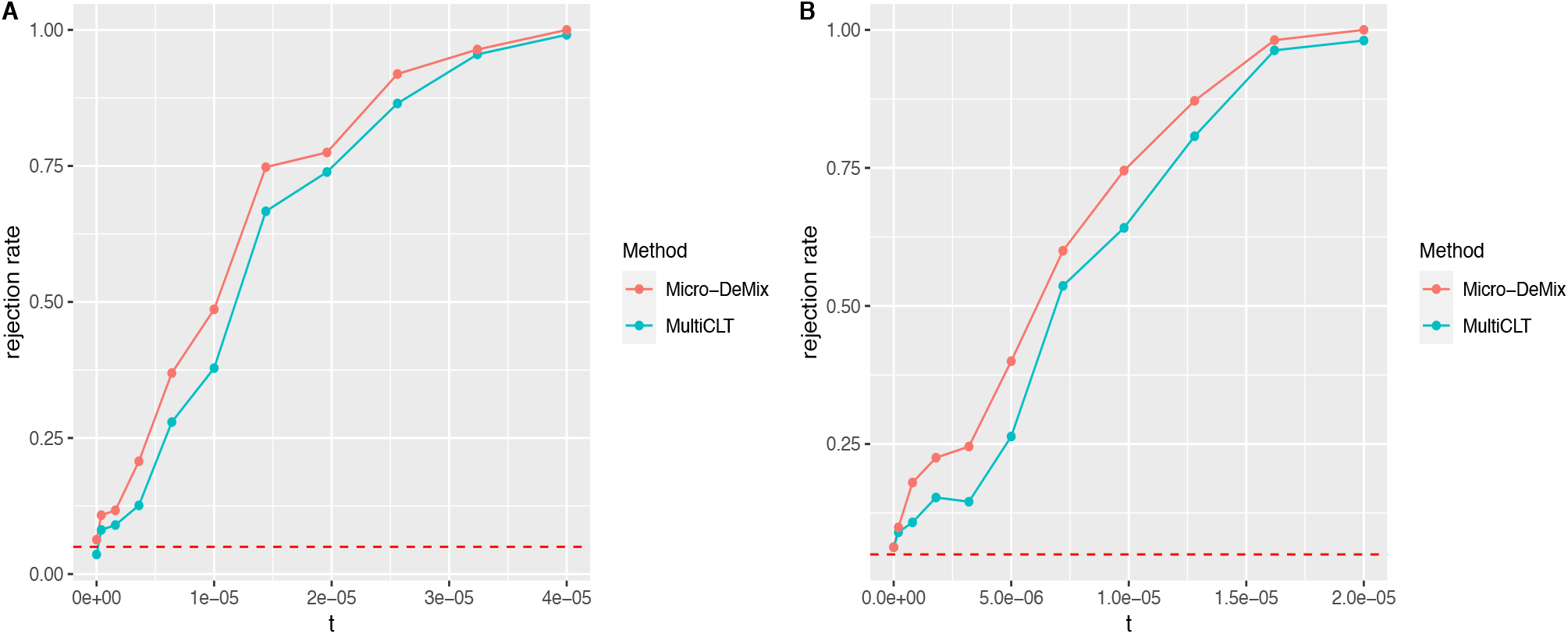
Type I error rates and powers obtained at various signal strengths from the simulation study. Plot A demonstrates power curves for G=10, while plot B demonstrates power curves for G=20. A horizontal dashed line is shown at 0.05. t on the x-axis denotes the total signal. Rejection rates were obtained using Micro-Demix (shown in red) and MultiCLT (shown in green).

In both settings, the power tends to increase as the total signal increases. Notably, Micro-Demix method outperforms MultiCLT in terms of power. This observation reveals that while multiCLT is valid under *H*_0_, it disregards the mixture of stool microbes, leading to lower power when *H*_0_ does not hold. The comparison demonstrates the importance of accounting for the sample mixture in the stool samples and the effectiveness of the proposed beta-multinomial model.

## 4 Application to the iHMP IBD Data

In this section, we revisit the integrative human microbiome project (iHMP) Inflammatory Bowel Disease (IBD) cohort, which is briefly discussed in the Introduction, to demonstrate the effectiveness and efficiency of the proposed Micro-DeMix in elucidating the composition of real fecal microbiome data. We are particularly interested in understanding how the fecal microbiome differs from the rectum microbiome in IBD populations, and how that difference may be related to the pathology of IBD.

As discussed in the Introduction, the IBD cohort was a longitudinal study, collecting taxonomic data on the microbiome from 132 individuals (104 IBD patients) by analyzing 1,785 stool samples and 651 intestinal biopsies over the course of one year [15]. This IBD cohort was part of the integrative human microbiome project (iHMP), which was designed to explore host–microbiome interplay to gain a holistic view of host–microbe interactions [15]. The Inflammatory Bowel Disease (IBD) Multi’omics Data (IBDMDB) research team collected microbiome data from different sites within the human gut of individuals with IBD to understand how the microbiome impacts human health and disease. For a fair comparison, our analysis focused on stool and rectum samples collected at the baseline, resulting in 13 samples for stool and 38 samples for rectum data in IBD patients. After excluding subjects with missing covariates of interest, such as age and gender, the stool group was narrowed down to 12 subjects. The OTU-level data for both groups are highly zero inflated. More than 90% of the entries in the OTU table derived from the stool samples are zero. Therefore, in the results below, we investigated the microbial composition of the fecal microbiome as compared to the rectum microbiome at higher taxonomic levels to mitigate the potential influences of excess zeros.

### 4.1 Phylum-level Analysis

We aggregated the absolute counts from the OTU level to the phylum level, resulting in 8 phyla, resulting a much lower percentage of zeros compared to the OTU-level data (50.9% vs. 93.6%). We noticed significant variations in total reads (library size) across samples for both groups, specifically, ranging from 1499 to 79228 for stool samples and from 34 to 31781 for rectum samples. We mitigated this issue by performing rarefying, using the R function rarefy even depth from package phyloseq as in the exiting literature [25]. In microbial studies, rarefying has been used for normalization by randomly subsampling the data to mitigate the potential influence of varying library sizes [20, 27, 16]. The rarefying procedure removes samples from the OTU table whose library sizes do not meet the user-defined rarefaction level (total counts) [27]. A reasonable rarefaction level in our analysis should maximize the inclusion of samples while removing samples with extremely low library sizes. We used 1499 as the rarefaction level for stool samples, as it is the minimum library size in stool group. For a fair comparison, we applied the same rarefaction level to rectum samples, resulting in 30 samples in rectum group. Since this procedure is random subsampling without replacement, we repeated the procedure 100 times and took the average to reduce variability for both groups.

We computed the relative abundance of these 8 phyla in both the rectum and stool samples, and the results revealed a clear difference in the microbiome composition (Fig. 2). Phylum Firmicutes was highly abundant in both the rectum and stool. It was the major phylum detected, representing more than 50% of the total microbial relative abundance in both groups. Overall, the relative abundance of Firmicutes in the stool (∼ 61.4%) was slightly higher than that in the rectum (∼ 52.0%). When compared to the microbial profile in rectum, the stool samples exhibited a considerable increase in the relative abundance of phylum Proteobacteria (from 7.1% to 30.1%), while the relative abundance of Bacteroidetes decreased from 38.1% to 6.7%. In addition, we observed an elevated relative abundance of the Verrucomicrobia phylum in the stool microbiome compared to the rectum, with an increase from 0.31% to 1.5%. Conversely, the phylum Fusobacteria showed a decrease from over 1.9% to a lower level. The remaining three phyla were nearly imperceptible in rectum and stool.

**Figure 2:**
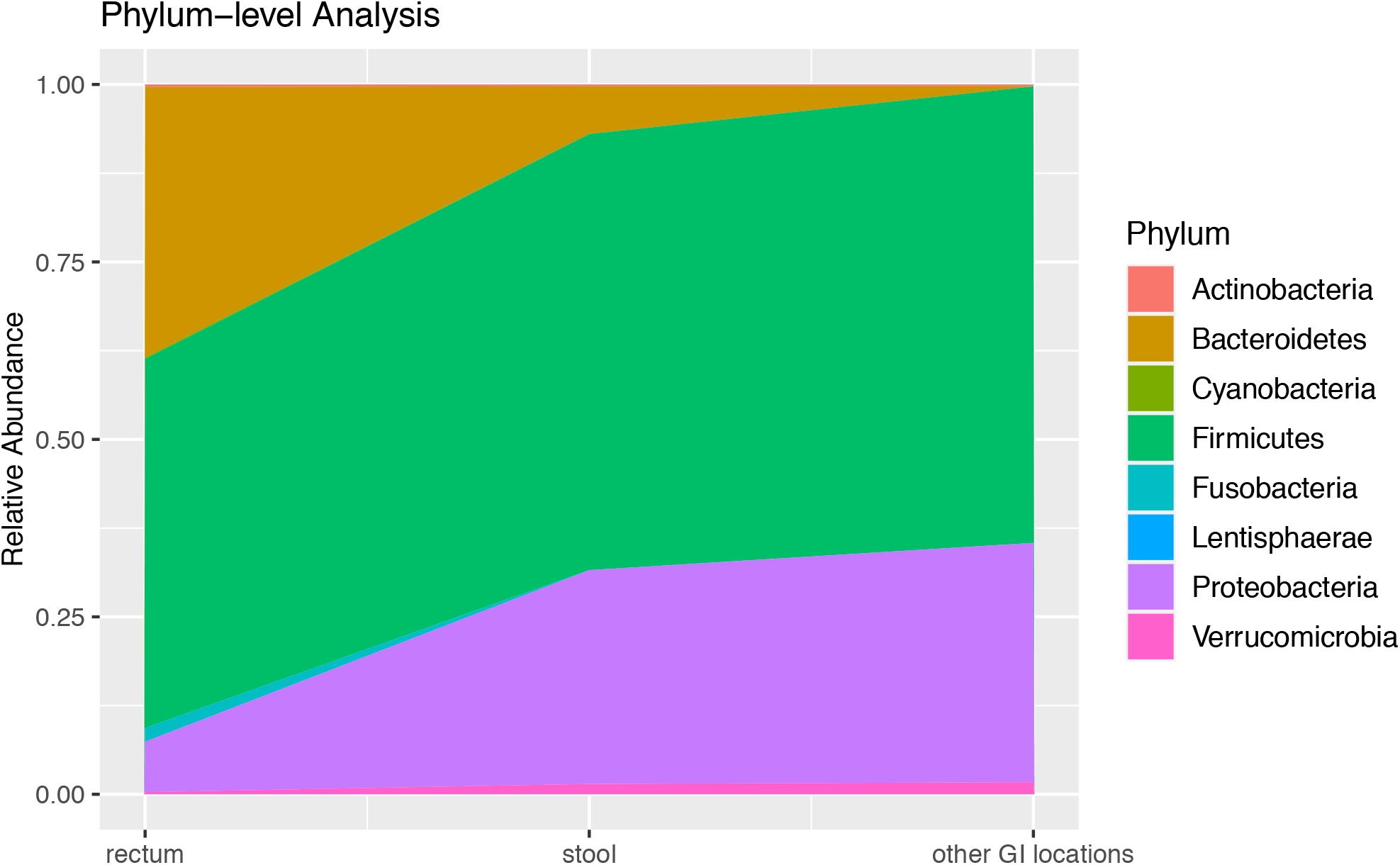
Relative abundance of microbial populations at the phylum level in the rectum, stool, and other GI locations.

To further understand the composition of the fecal microbiome in IBD populations, we fitted the proposed Micro-DeMix model, incorporating gender and age as covariates. This analysis revealed estimates of the relative abundance of selected phyla from the other GI locations (Figure 2). Compared to the rectum microbiome, the microbial profile of other GI locations revealed a higher relative abundance of phylum Firmicutes (∼ 64.3%), Proteobacteria (∼ 33.7%) and Verrucomicrobia (∼ 1.7%). Phylum Bacteroidetes was undetectable in other GI locations. The remaining phyla also exhibited relatively low abundance (*<* 1%). Based on 10,000 simulated data sets, the Micro-DeMix test suggested significant differences in the taxonomic composition of the 8 phyla between the rectum and other GI locations in IBD populations (p-value *<* 1*e* 4).

Our findings are consistent with recent studies on the gut microbiome in IBD populations. Specifically, the rectum and fecal microbiome of IBD patients is dominated by three major bacterial phyla: Firmicutes, Bacteroidetes, Proteobacteria, and to a lesser degree Verrucomicrobia and Actinobacteria [11, 10]. Our results are also consistent with recent studies that described distinct changes in the composition of the microbiota along the gastrointestinal tract in IBD populations. The lower gastrointestinal tract exhibited a larger abundance of Firmicutes and Bacteroidetes, whereas the upper gastrointestinal tract was predominated by Proteobacteria and Firmicutes [12]. Compared to healthy individuals, the reduction in the relative abundance of Firmicutes and an elevation in Bacteroidetes have been observed in IBD patients [11, 12, 13, 19]. Thus, our results shed light on the scientific premise of developing an effective microbiome-based treatment approach for IBD, such as the therapeutic supplementation of probiotics, prebiotics, and synbiotics, and fecal microbiota transplantation (FMT).

### 4.2 Microbial Composition within Proteobacteria and Firmicutes

In Section 4.1, we identified Proteobacteria and Firmicutes as the most abundant phyla in stool samples. To further understand the microbes in these two phyla, we conducted analyses of microbial composition at lower taxonomic levels (class, order, family) within Proteobacteria and Firmicutes phylum (Fig 3).

**Figure 3:**
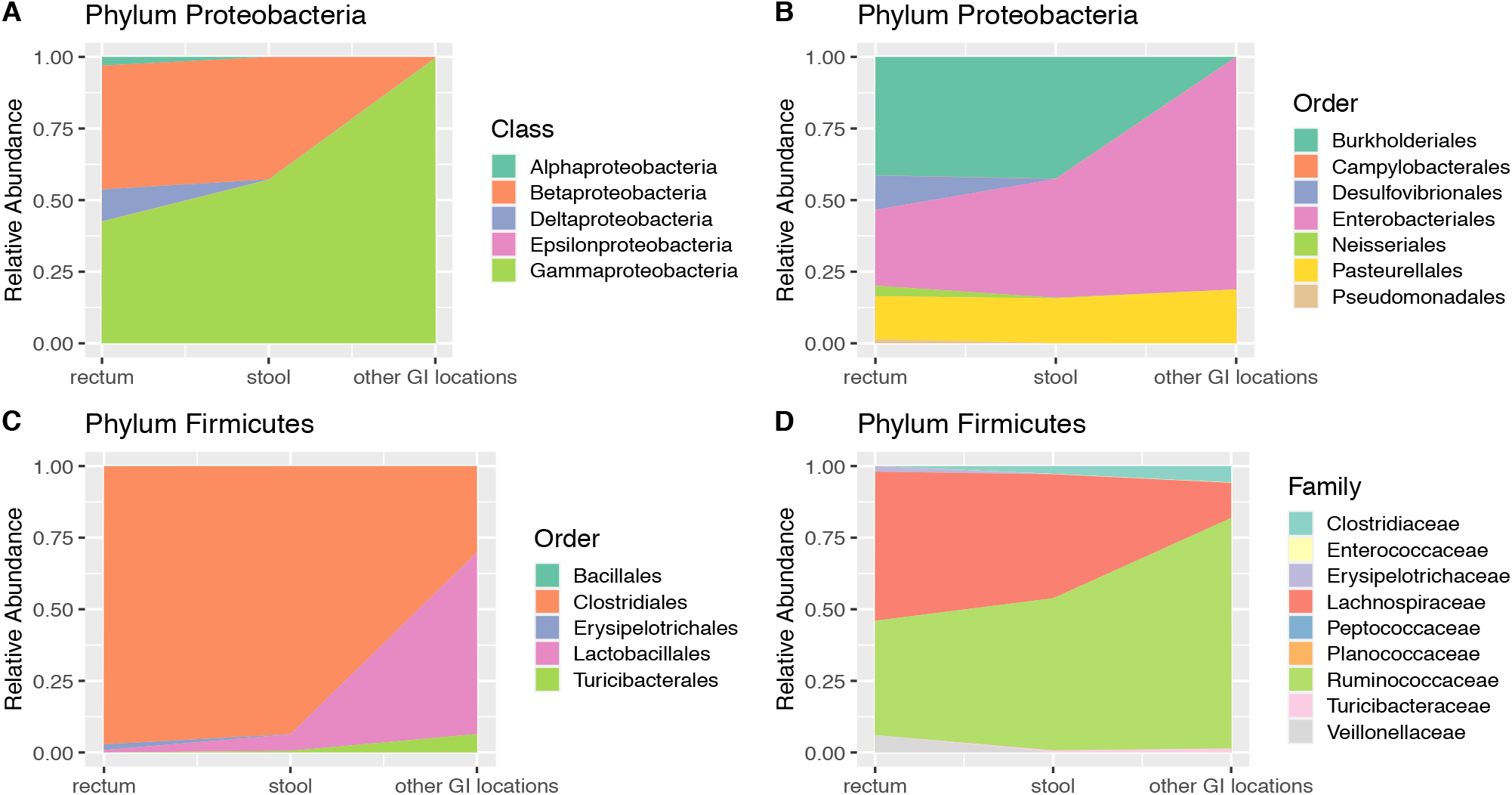
Relative abundance of microbial populations at various taxonomic levels in the rectum, stool, and other GI locations. Plot A and B demonstrate the microbial composition at the class and order levels within the phylum Proteobacteria. Plot C and D demonstrate the microbial composition at the order and family levels within the phylum Firmicutes.

Similar to the analyses in Section 4.1, we computed the relative abundance of all microbial classes and orders within phylum Proteobacteria for both the stool and rectum samples, and implemented the Micro-DeMix model to elucidate the difference between the rectum microbiome and the microbes in other GI locations. We followed the rarefying procedure that described in Section 4.1 to process the stool OTU data. In stool samples, the minimum library size is 13, which is not big enough to be included in the study. Therefore, we used 369, the second minimum, as the rarefaction level for both stool and rectum group. The outcomes of our analyses are visually represented in Figure 3A and Figure 3B.

Figure 3A reveals that within phylum Proteobacteria, the major classes in rectum and stool are Betaproteobacteria and Gammaproteobacteria. However, the rectum microbiome is distinguished by a higher relative abundance of class Deltaproteobacteria (∼ 11.1%) and Alphaproteobacteria (∼ 3.0%), which are significantly low in abundance in stool samples. There is a low relative abundance of Epsilonproteobacteria (*<* 1%) in both stool and rectum. As seen in Fig 3A, Gammaproteobacteria (∼ 99.6%) is the most dominant class among the selected classes in other GI locations, while the other 4 classes are nearly undetectable. Although this finding may appear extreme, it is aligned with the existing knowledge that IBD patients have an altered gut microbiota marked by an increased relative abundance of Gammaproteobacteria class [29].

We proceeded to analyze the microbial composition of the phylum Proteobacteria at the order level, revealing marked differences between the rectum and stool samples (Figure 3B). While both showed high abundance in orders Burkholderiales, Enterobacteriales, and Pasteurellales, the rectum samples was mainly charaterized by the orders Desulfovibrionales and Neisseriales. We utilized the Micro-DeMix method to assess the microbiome composition in other GI locations. Our analysis revealed a dominance of Enterobacteriales (∼ 81.1%), along with a lower proportion of Pasteurellales (∼ 18.83%). This finding is supported by a recent study showing that Enterobacteriaceae exhibit increased abundance in the low-pH GI location and ileum [26].

The Micro-Demix test indicates significant differences in microbial profiles between the rectum and other GI locations (p-value *<* 1*e* − 4) within the phylum Proteobacteria at both the class and order levels.

We further explored the microbial composition of the Firmicutes phylum. In the stool samples, three classes were identified: Bacilli, Clostridia, and Erysipelotrichi. At the original OTU-level, class Clostridia dominated with a percentage of 97.1%, while class Erysipelotrichi was nearly undetectable. Given these findings, we narrowed our focus to lower taxonomic levels, specifically, order and family within the Firmicutes phylum. Similarly, we employed the minimum library size, 493, as the rarefaction level and processed both stool and rectum OTU data. The corresponding results are shown in Figure 3C and Figure 3D.

Notably, at the order level (Figure 3C), the phylum Firmicutes in the rectum and stool was predominantly represented by the order Clostridiales (97.1% in rectum and 93.6% in stool of all Firmicutes). Our results demonstrate the presence of order Turicibacterales in the stool samples, while it is absent in the rectum samples, which suggest that order Turicibacterales in stool samples may originate from other GI locations. The relative abundance of the order Lactobacillales increased in stool compared to rectum from 0.86% to 5.7%. In contrast, Order Erysipelotrichales is found to be present (2.0%) in the rectum but remains at lower levels in the stool. The results of Micro-DeMix analysis show a p-value *<* 1*e* − 4 for testing differential abundance between the rectum and other GI locations. Meanwhile, it highlights a significantly higher relative abundance of Lactobacillales (∼ 63.4%) in other GI locations compared to the rectum and stool. The analysis estimates that order Turicibacterales (∼ 6.4%) is also more abundant in other GI locations. These findings indicate a distinct microbial composition in other GI locations, with increased prevalence of order Lactobacillales and Turicibacterales.

At the family level within the Firmicutes phylum (Figure 3D), major families of the rectum microbiota are Ruminococcaceae, Lachnospiraceae, and Veillonellaceae. Comparisons between stool and rectum samples reveal an increase in Ruminococcaceae and a decrease in Lachnospiraceae and Veillonellaceae, aligning with findings from other studies [11]. One significant finding is the presence of the Clostridiaceae, Enterococcaceae, Planococcaceae, and Turicibacteraceae families in stool samples but not in the rectum. This observation suggests that these families detected in stool originate from other gastrointestinal (GI) microbiome community.

In other GI locations, our analysis shows a higher abundance of the Clostridiaceae and Ruminococcaceae families. In contrast, the family Lachnospiraceae exhibits a comparatively lower abundance. Furthermore, the p-value obtained through Micro-Demix hypothesis testing is low (*<* 1*e* − 4). These findings imply distinctive microbial profiles among the rectum, stool, and other GI locations, indicating variations in the relative abundance of specific taxa.

## 5 Conclusion

In this paper, we introduced Micro-Demix, a beta-multinomial model designed for deconvoluting microbial abundance in stool samples. Additionally, we developed a hypothesis testing framework for detecting differential abundance, which has proven more effective compared to existing methods. We applied Micro-Demix to rectum and stool microbiome datasets from the iHMP IBD cohort in elucidating the composition of real fecal microbiome data in IBD populations.

In the analyses of the iHMP IBD data, we performed rarefying on stool and rectum datasets to address the large variations in library sizes across samples. While rarefaction has been widely adopted in real microbiome applications, a limitation of this approach is that the selection of rarefaction level is data-dependent, and different rarefaction levels may lead to slightly different results. Due to the limited sample size, our strategy for selecting the rarefaction level is to keep as many samples as possible while removing samples with unreliably low library sizes. Different strategies may be considered for other data sets.

As mentioned earlier, the current paper focuses on integrating rectum samples and fecal samples due to data availability and biological relevance. However, the proposed method can seamlessly be used for integration of fecal samples with microbiome data collected from other GI locations. Furthermore, with microbiome data available from two or more GI locations, we could extend Micro-DeMix to have a refined decomposition of the fecal microbiome. Specifically, following work could consider the stool microbiome as a *J*-component mixture of gut microbes, i.e., we let 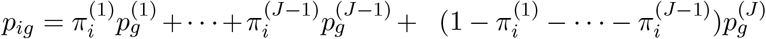 in (1), where ***π***_*i*_ follow a Dirichlet(***a***_*i*_) distribution. We leave this fruitful area for future research.

## 6 Appendix

### 6.1 Mean and Variance of *y*_*ig*_

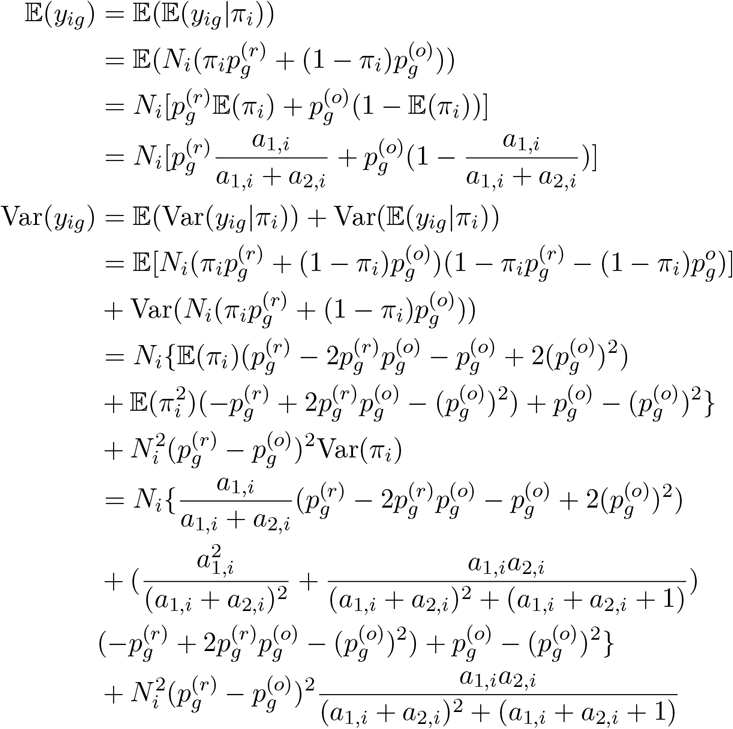

### 6.2 Proof of Proposition 1

Since *N* ^(*r*)^*/N → K >* 0, and noting that **y**_*i*_ and 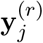 are independent, by the central limit theorem, we have

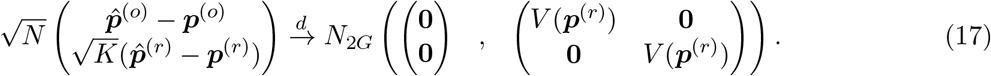

Let 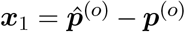, 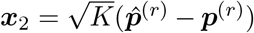, and let (***y***_1_, ***y***_2_) be a random vector with the multivariate normal distribution in (17). Then we can write (17) as

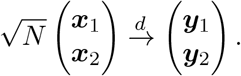

Define a continuous function 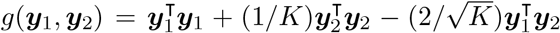. It is seen that 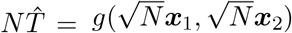. By the continuous mapping theorem, we obtain 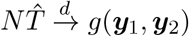.

## 7 Data Availability Statement

The IBD data analyzed in the paper can be accessed via HMP portal at https://portal.hmpdacc.org/. Code for implementing our methods is available in the R package MicroDemix, available at https://github.com/liuruoqian/MicroDemix.

## 9 Author contributions statement

R.L., Y.W. and D.C. conceived and conducted the experiments. R.L., Y.W. and D.C. analysed the results, wrote and reviewed the manuscript.

## 10 Acknowledgments

Yue Wang is supported in part by the funding R01GM145772 for the National Institute of Health. Dan Cheng is supported by NSF Grant DMS-2220523 and Simons Foundation Collaboration Grant 854127.

